# Phase-Dependent EEG Decoding of Continuous Visual Stimuli

**DOI:** 10.1101/2025.11.10.687532

**Authors:** Michele Deodato, David Melcher

## Abstract

Neural oscillations support cognition and perception by organizing information across oscillatory cycles, such that different phases correspond to distinct computational states. Although animal studies strongly support this view, evidence in humans is mixed and largely based on brief, flashed stimuli. Here, we tested whether alpha oscillations, the dominant human brain rhythm, rhythmically modulate visual encoding during sustained perception. EEG was recorded while participants viewed long-lasting (2s) Gabor patches. Using a phase-dependent decoding approach, we extracted EEG activity at specific alpha phases and quantified stimulus decoding accuracy. Decoding performance varied systematically with alpha phase, following a smooth unimodal profile that peaked over occipito-parietal electrodes. These effects emerged well after stimulus onset, indicating a sustained modulation of visual processing during continuous input. Our findings demonstrate that visual information is encoded more accurately at specific alpha phases. This provides direct evidence that sensory processing is gated by intrinsic brain activity and introduces a generalizable decoding framework for linking oscillatory phase to neural information processing.

## Introduction

It is widely accepted that our experience of visual space is actively constructed by the brain. Despite sharp color vision being limited to the fovea and blind spots lacking any input, we perceive a seamless and detailed visual field. Likewise, while we experience vision as continuous, many temporal aspects of sensory processing may be influenced by spontaneous rhythmic patterns of neural activity (1, 2). According to temporal sampling hypotheses, the processing of incoming information is shaped by neural oscillations in the alpha frequency range (8-13 Hz) (2). These rhythmic fluctuations in neural excitability act as dynamic gates, modulating neural firing and thus creating temporal windows during which sensory input is preferentially sampled (3, 4). By segmenting sensory processing, the visual system optimizes the extraction of relevant information from a continuous sensory stream under constraints due to eye movements and limits in attentional and neural processing capacity.

In support of this account, several studies have shown that the phase of pre-stimulus alpha oscillations influences the perception of phosphenes (5), illusory flashes (6), and near-threshold flashes (7), as well as temporal integration (8, 9) and order and timing judgments (10, 11) of brief visual events. Additionally, intra- and inter-individual differences in the frequency of alpha oscillations have been linked to the temporal resolution of perception (12–14). However, findings are mixed (15), with recent work challenging the reliability and generalizability of these effects (16, 17). More critically, three major limitations have hindered a deeper understanding of the relationship between oscillations and sensory processing. First, effects are largely confined to transient stimuli, limiting their explanatory power for continuous, naturalistic visual processing. Second, research has focused almost exclusively on pre-stimulus phase, leaving open the question of whether sensory encoding is dynamically modulated during sustained visual input. Finally, rhythmic processing is typically assessed via subjective reports rather than neural measures of information processing, potentially being influenced by decision making factors.

Here, we address these gaps by shifting focus from perceptual reports of brief flashes to neural signal dynamics of continuous stimuli. We recorded EEG from participants viewing long-lasting (2 s) Gabor patches while performing an unrelated task designed to maintain fixation and stable visual input (Fig. 1). The Gabor stimuli could vary in orientation (-45 and 45 degrees) and spatial frequency (1.85 and 7.5 cpd) and were entirely task-irrelevant, allowing us to examine visual processing in the absence of attention or behavioral relevance. The functional significance of neural oscillations rests on the idea that different phases of the cycle correspond to distinct computational states, a notion well supported by animal studies (4, 18). To test that neural processing of continuous stimuli fluctuates rhythmically, we developed a phase-dependent decoding approach that quantifies how accurately visual information can be read from neural signals as a function of the phase of ongoing oscillations. Rather than making claims about conscious experience, our analyses quantify how the decodability of visual stimuli from EEG fluctuates as a function of oscillatory phase. This approach provides a direct measure of the information content in neural signals, rather than subjective perception per se. Our results demonstrate that neural decoding of continuous visual input varies unimodally across the alpha cycle, providing direct evidence that visual processing is rhythmically modulated in time, in synchrony with brain oscillations.

**Figure 1.**
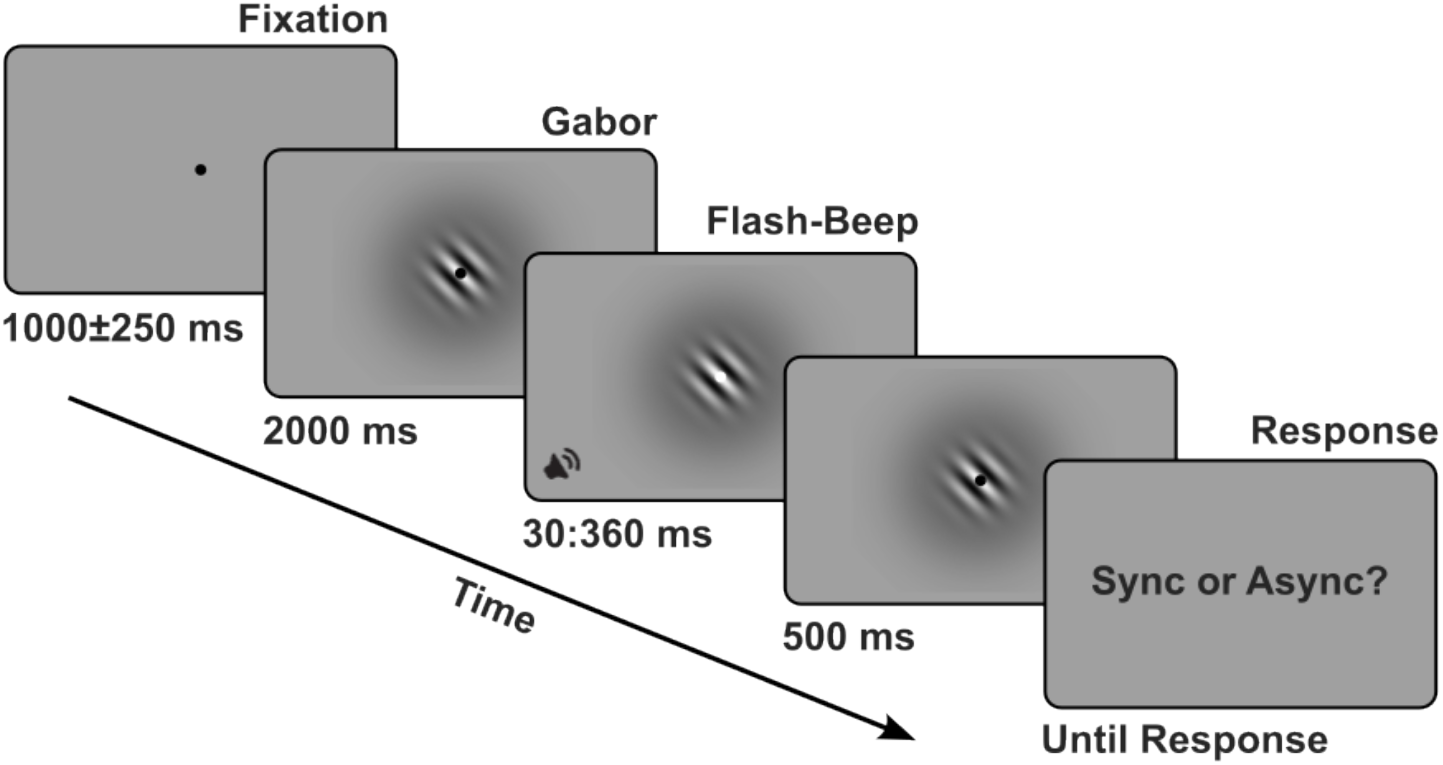
Experimental Design. Schematic of an example trial. Each trial began with a central fixation cross, followed by the onset of a centrally presented Gabor patch, which remained onscreen for a prolonged duration of 2000 ms. At the end of this period, participants were presented with an audiovisual pair (30 ms flash and 30 ms tone) with stimulus onset asynchronies (SOAs) and order varying randomly across trials. After a 500 ms delay, participants responded whether the flash and beep appeared synchronous or asynchronous.

## Results

### Distinct temporal stages in EEG representations of continuous stimuli

To quantify neural encoding of stimulus features, we applied time-resolved multivariate pattern analysis, training a linear discriminant analysis (LDA) classifier to decode orientation and spatial frequency from EEG topographies at each time point (–500 to 2000 ms relative to stimulus onset).

Decoding accuracy for stimulus orientation remained at chance level, likely due to the oblique orientation angles (19), stimulus size and the fine-scale cortical encoding of orientation, which is difficult to resolve with scalp EEG-based LDA. In contrast, spatial frequency could be reliably decoded from EEG signals. Decoding accuracy rose significantly above chance shortly after stimulus onset (p_cluster_ < 0.05), peaked around 120 ms post-stimulus (mean peak accuracy = 91.45%, SD = .07), and remained significantly above chance throughout the 2-second presentation (Fig. 2).

**Figure 2.**
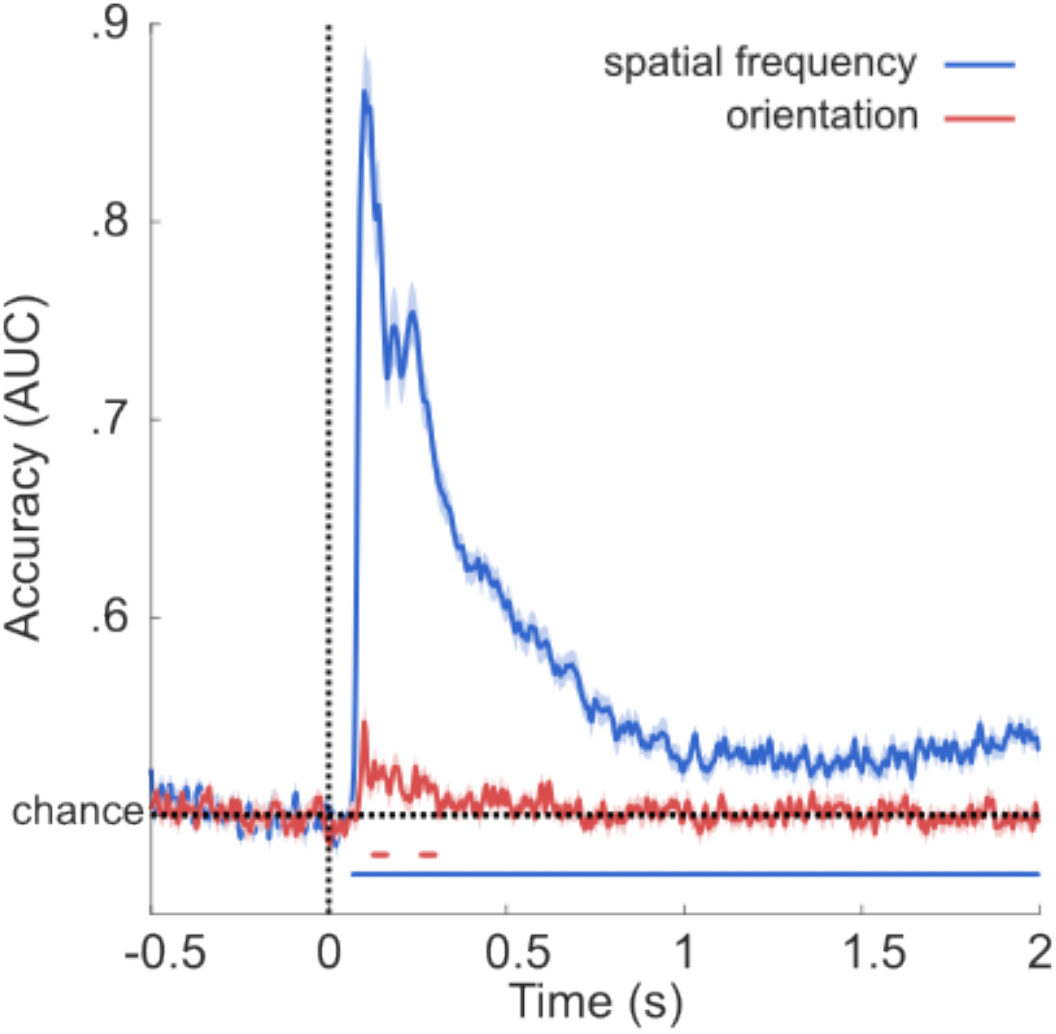
EEG Decoding time-locked to stimulus onset. Decoding accuracy for stimulus orientation (red) and spatial frequency (blue) over time, locked to stimulus onset (time = 0). Orientation decoding showed brief, weak above-chance performance shortly after stimulus onset but remained at chance for the rest of the trial. Spatial frequency decoding shows two distinct stages: an early encoding stage (0–500 ms) marked by high accuracy, and a later maintenance stage (1000–2000 ms) during which decoding declined but remained reliable. Shaded areas indicate ±1 SEM across participants. Horizontal lines indicate significant (p<.05) decoding accuracy.

Notably, decoding performance showed an exponential decay characterized by an early encoding stage (0–500 ms) with transient peaks of high accuracy, followed by a later maintenance stage (1000–2000 ms) during which accuracy declined (<60%) but remained above chance. Previous research has linked changes in neural activity, in particular in alpha band power, to microsaccades (20). To verify that this residual decoding performance was not driven by ocular artifacts, we conducted two control analyses: decoding using only horizontal and vertical eye-position data, and a comparison of microsaccade rates between low and high spatial frequency trials. Both analyses yielded no significant results, confirming that the observed EEG effects were not driven by eye movements. This suggests that while spatial frequency information is strongly encoded immediately after stimulus onset, it continues to be represented in a sustained format during prolonged visual stimulation. These results establish that continuous, unattended visual stimuli elicit robust and temporally extended neural responses that can be tracked using EEG.

### Phase-dependent decoding reveals rhythmic gating of visual information

To test whether neural processing of continuous visual input fluctuates rhythmically with alpha phase, we performed decoding analyses on EEG data selectively sampled at distinct points within the alpha cycle. We restricted this analysis to a late time window, between 1000 and 2000 ms after stimulus onset, when the stimulus was still present but transient onset-related activity had long subsided. This choice is essential, since onset responses elicit pronounced alpha desynchronization that compromises a precise and meaningful estimation of alpha phase (see methods).

Because alpha phase varies across the scalp, we began by computing the instantaneous phase of alpha oscillations (8–13 Hz) from a single electrode. For each trial, we identified time points corresponding to the peaks and troughs of the alpha cycle at that electrode and extracted the corresponding scalp voltage topographies (Fig. 3A). We then computed decoding accuracy for spatial frequency separately for peak and trough phases. A two-tailed paired t-test was used to compare accuracy between the two phases (Fig. 3B).

**Figure 3.**
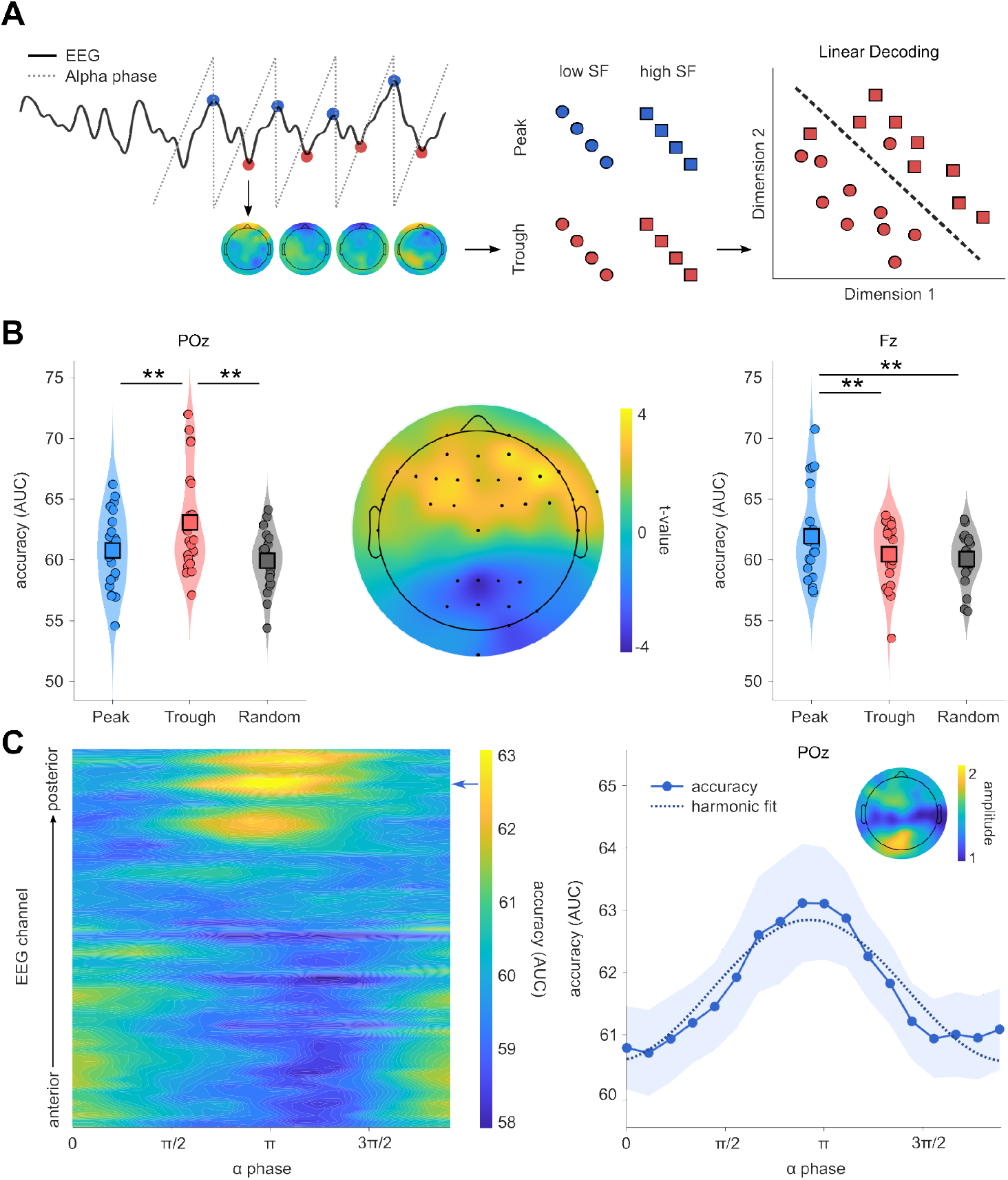
Phase-dependent EEG Decoding. (A) Schematic of the phase-dependent decoding approach. For each trial, scalp voltage topographies were extracted at specific points of the alpha cycle (e.g., troughs) and used to train a linear decoder to classify stimulus identity. (B) Decoding accuracy was compared between alpha peaks and troughs. The violin plots show that posterior channels (e.g., POz) showed significantly higher decoding accuracy at the trough, while frontal channels (e.g., Fz) showed higher accuracy at the peak. The topographic map displays the t-values from paired comparisons across electrodes (peak vs. trough), with significant clusters identified via permutation testing. Accuracy at both sites exceeded that observed when phase samples were selected randomly. (C) The color map shows tuning of decoding accuracy as a function of alpha phase (18 bins) and scalp location (64 channels). The plot on the right side shows the tuning curve corresponding to one row of the map (blue arrow, channel POz) with the corresponding harmonic regression fit (dotted line). Shaded area is the SEM across participants. The topographic map shows the amplitude of the fitted sinusoid, reflecting the strength of phase-dependent modulation across the scalp and peaking over posterior electrodes.

This procedure was then extended to all electrodes and resulting t-values were subjected to a cluster-based permutation test. Two significant clusters emerged (p_cluster_ < 0.05): one over occipito-parietal electrodes showing higher decoding accuracy at the trough, and another over frontal electrodes exhibiting higher accuracy at the peak (Fig. 3B). This pattern remained robust with respect to a control analysis using randomly sampled phases, confirming that the effect was phase-specific (Fig. 3B). Additionally, the effect disappeared when considering the phase of theta (4-7 Hz) oscillations, showing that the effect is also frequency specific. These findings demonstrate that the strength of neural representations is not constant over time, even when visual input remains unchanged, but instead varies systematically with the phase of ongoing alpha oscillations, showing that visual processing is shaped by the intrinsic temporal structure of ongoing neural activity.

Using 18 equally spaced phase bins spanning 0–2*π*, decoding accuracy formed smooth unimodal tuning curves (Fig. 3C). A harmonic regression model confirmed the significance of this rhythmic modulation. Specifically, the model explained a substantial portion of variance in decoding accuracy across the whole scalp (mean R^2^ =.57, range = [.47, .69], p_cluster_ < 0.001), with a mean modulation amplitude of 1.5% (SD =.22%) (Fig. 3C). Preferred phases were consistent across participants within occipital and frontal clusters (Rayleigh test p < 0.05), although this phase alignment did not survive correction for multiple comparisons (see methods). This variability is expected as it reflects individual differences in cortical geometry, the location and orientation of cortical alpha generators. This analysis confirm that decoding accuracy is rhythmically modulated by ongoing alpha phase, with a single optimal phase for visual processing, consistent with a rhythmic gating mechanism.

### Phase-dependent decoding depends on anti-phase signals

The presence of opposite phase-dependent decoding effects at frontal and posterior sites raises the question of whether these patterns reflect two interacting sources. While conclusions from scalp EEG data can only be tentative, to address this question we estimated the phase relationship between each pair of significant anterior–posterior EEG channels by computing cross-spectral density in the alpha band.

Circular V-tests showed that 215 out of 243 anterior–posterior channel pairs (88.5%) exhibited significant clustering around 180° (p_Bonferroni_ < 0.05; mean phase offset = 179.34°) (Fig. 4). Additionally, the Phase-Slope Index, a measure of information flow robust to volume conduction (21), between frontal and posterior channels was not different from 0 (p_permutations_ > 0.05), suggesting no interaction between these regions (Fig. 4).

**Figure 4.**
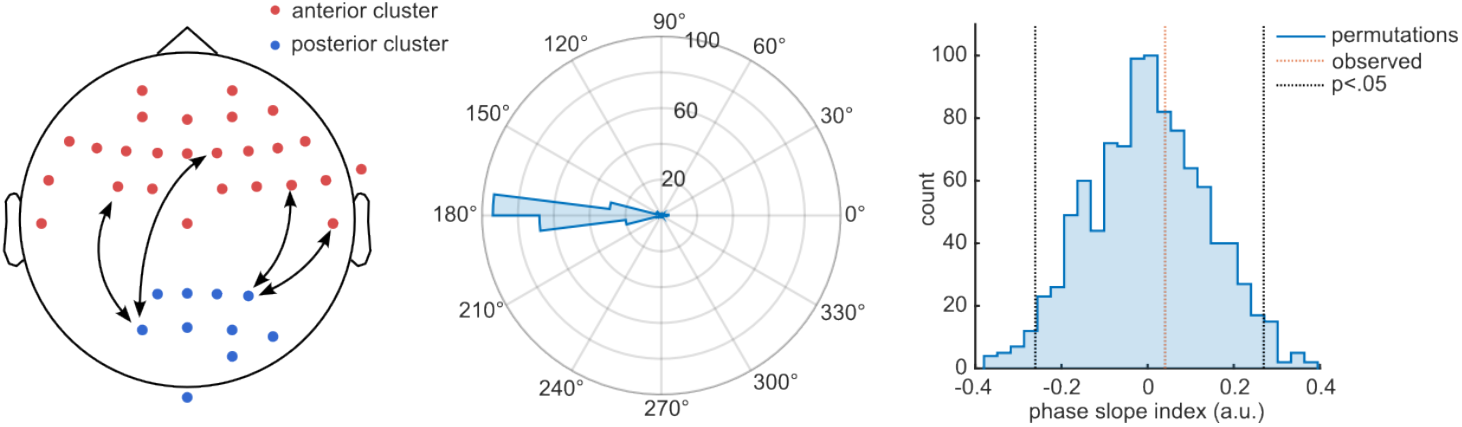
Distribution of alpha phase differences between anterior and posterior clusters. (Left) Scalp map showing significant frontal (red) and occipito-parietal (blue) channels identified in the phase-dependent (peak vs trough) decoding analysis. Black arrows indicate anterior–posterior channel pairs. For each possible pair, we computed the average phase difference in the alpha frequency range (8–13 Hz) and the phase slope index (PSI). (Middle) Polar histogram showing the distribution of alpha phase differences across all channel pairs (N=243) averaged across participants. The distribution is tightly clustered around 180°. (Right) Random distribution of the PSI. The observed PSI (averaged across anterior-posteior pairs, red dotted line) was not significant with respect to the upper and lower distribution limits (black dotted lines), indicating no evidence for information transfer between channels.

These results do not support interacting sources in distal areas but are consistent with previous work identifying posterior alpha generators, likely within the occipito-parietal cortex, as a dominant generator of rhythmic visual processing (8, 22). However, given the limited spatial resolution and inherent volume conduction in sensor-level EEG data, we cannot exclude the presence of multiple interacting sources within these areas or more complex network dynamics contributing to the observed phase patterns.

## Discussion

Oscillatory brain activity is a core organizing principle of neural function and a well-established marker of dysfunction in a range of clinical conditions (23). Understanding how these rhythms influence sensory processing is therefore central to neuroscience. Here, we provide direct evidence for a rhythmic modulation of visual processing during sustained visual stimulation. Specifically, by showing that decoding of visual information fluctuates with ongoing alpha phase, we demonstrate that rhythmic influences on sensory processing persist throughout continuous visual input. This provides a strong neural foundation for the idea that perception is shaped not only by what is seen, but also by when it is seen within the brain’s intrinsic temporal structure.

Much of the existing research on phase-dependent visual processing has relied on participants’ subjective reports of bistable stimuli (6, 8, 24), leaving open questions about whether these rhythmic fluctuations also shape objective neural representations. While some studies have shown that pre-stimulus alpha phase can modulate early sensory responses, such as evoked potentials and global field power (25–27), these effects reflect the initial encoding of stimuli. In contrast, our analyses reveal that alpha phase continues to modulate neural representations well beyond stimulus onset, during sustained visual processing. This distinction is critical, as our results highlight meaningful differences between initial encoding and maintenance stages of processing. We reveal a persistent, spontaneous oscillatory influence on neural processing that would be masked in more conventional paradigms. Our focus on continuous visual input aligns with growing efforts to shift experimental paradigms toward the temporal dynamics of natural vision (28, 29). Furthermore, because stimulus onset can elicit phase resetting (30) and feature-specific changes in fixational eye-movements (31), isolating the maintenance period allowed us to reveal ongoing, spontaneous rhythmic influences on neural processing that would otherwise be obscured. Notably, the poor replicability of some of the original findings may stem from the influence of stimulus onset and offset responses, the artificial and technically demanding setups required to precisely control millisecond-scale transients and the participants effort (or lack thereof) in detecting subtle differences, rather than from a lack of underlying rhythmicity. By demonstrating phase-dependent neural decoding during continuous viewing, we provided a robust and more ecologically valid test of the relationship between neural oscillations and visual processing.

Over the past century, different interpretations of this relationship have gained traction, often motivated by compelling psychological phenomena (e.g., (32). Yet enthusiasm has repeatedly waned due to the lack of definitive neural evidence (1, 2). One of the earliest formalizations of this idea came from Pitts and McCulloch, who proposed a ‘cortical scanning’ theory in which cortical layers cyclically fluctuate in receptivity to incoming input (33). More recently, the framework of rhythmic perception proposed that perception is not uniformly continuous, but modulated by the phase of ongoing brain rhythms such that certain phases promote more efficient sensory processing, while opposite phases impair it (3). Importantly, and consistent with our findings, this rhythmicity does not imply discrete, frame-like snapshots of perception, but rather continuous fluctuations in neural excitability that modulate the fidelity of sensory encoding over time.

Rhythmic fluctuations are often attributed to attentional sampling, rather than intrinsic periodicity within sensory processing per se (10, 32, 34, 35). In this context, oscillatory effects arise from rhythmic modulations of sensory gain driven by fluctuations in attentional focus. However, growing evidence indicates that attentional and sensory rhythms coexist and operate at distinct frequencies, with attention often linked to slower theta rhythms (4-7 Hz) (36), and early sensory encoding associated with faster alpha-band oscillations (8-13 Hz) (37). This supports the idea that multiple, coexisting oscillatory mechanisms coordinate different aspects of perception and cognition at different temporal scales (8, 37, 38).

In the present study, the stimuli were entirely task-irrelevant, and participants were not instructed to attend to them. This makes it unlikely that fluctuations in decoding performance were driven by rhythmic attentional enhancement. Instead, these results may reflect a form of rhythmic suppression, where sustained attention to fixation periodically inhibits processing of irrelevant inputs, leading to phase-dependent modulations in visual processing (35, 36).

An alternative explanation that does not invoke attention is that alpha phase reflects the responsivity of spatial frequency–selective neural populations. For example, recent work suggest that alpha oscillations could reflect differential involvement of the magnocellular and parvocellular pathways (39, 40), which could explain why phasic effects are often observed with transient, low-contrast stimuli that predominantly engage the magnocellular pathway (7, 41). More generally, posterior alpha troughs have been associated with periods of heightened cortical excitability in visual areas, supporting enhanced sensory processing. Correspondingly, fluctuations in the balance between synaptic excitation and inhibition have been shown to modulate the strength of alpha-band rhythmic inhibition (42) and its impact on perception (43). Overall, these interpretations support the central conclusion that visual processing of continuous stimuli fluctuates rhythmically as a function of alpha phase.

On a final methodological note, our phase-dependent decoding technique offers a flexible and generalizable approach for probing how oscillatory dynamics shape cognitive processing across diverse experimental contexts. By isolating specific moments in the oscillatory cycle, such as peaks and troughs, we demonstrate that neural processing of visual input fluctuates systematically, revealing when decoding from EEG is most effective. Beyond its theoretical implications, this approach is especially promising for brain–computer interface (BCI) applications, where timing decoding to optimal oscillatory phases could significantly enhance performance. We showed that not all moments in the EEG signal are equally informative: decoding accuracy can vary markedly with ongoing oscillatory phase, offering a principled way to enhance predictions of perception from continuous EEG activity.

## Materials and Methods

### Participants

Twenty-two participants (12 females) between 18 and 35 years old (age mean: 23.13 years; SD: 3.6) participated in the experiment and received compensation. Inclusion criteria were normal or corrected to normal vision and English fluency. Data were collected in accordance with the Declaration of Helsinki, and the study protocol was approved by the local ethics committee (New York University Abu Dhabi IRB).

### Procedure

The experiment took place in a dark, magnetically shielded room. Stimuli were projected with a PROPixx projector (VPixx Technologies) set to a refresh rate of 100 Hz, displayed on a 65 cm × 36 cm screen. The experiment was implemented using MATLAB (The MathWorks) and the psychophysics toolbox (44). Participants laid their head on a chin rest at a distance of 90 cm. Eye data were recorded with an Eyelink 1000 eye-tracker (SR Research, Ontario, Canada) in desktop mount mode with a sampling rate of 1000 Hz.

Upon arrival to the laboratory, participants were instructed on the good practices to follow during EEG experiments (e.g., limit body movements and blinking) and were prepared for EEG recording. Each trial began with a central fixation cross presented for 1000 ms, with an added temporal jitter of ±250 ms. This was followed by the appearance of a centrally presented Gabor patch, which remained on screen for 2000 ms. The Gabor stimuli varied across trials in orientation (–45° or 45°) and spatial frequency (1.85 or 7.5 cycles per degree, cpd). At the end of the 2-second Gabor presentation, a brief stimulus pair was presented. This was either a 30 ms, 2000 Hz tone paired with a 30 ms white luminance flash, or two successive flashes, implemented by changing the fixation cross to white. The temporal order (flash-first or beep-first) and stimulus onset asynchrony (SOA) varied randomly across trials. For audiovisual pairs, SOAs ranged from 0 to 360 ms in 30 ms steps; for double-flash trials, SOAs ranged from 50 to 230 ms in 20 ms steps. The Gabor remained on screen for an additional 500 ms after the stimulus pair, after which participants reported whether the flash and beep were synchronous or asynchronous, or indicated the number of flashes perceived. Participants were instructed to blink only after the Gabor had disappeared. The experiment consisted of 7 blocks (100 trials each) with audiovisual stimulus pairs, followed by 3 blocks (107 trials each) of flash-flash trials. Short breaks were provided between blocks. The full session lasted approximately 2.5 hours.

### EEG recording and preprocessing

EEG data were recorded using a 64-channel active electrode system (BrainProducts GmbH) arranged according to the international 10–10 system (Fp1, Fz, F3, F7, FT9, FC5, FC1, C3, T7, TP9, CP5, CP1, Pz, P3, P7, O1, Oz, O2, P4, P8, TP10, CP6, CP2, Cz, C4, T8, FT10, FC6, FC2, F4, F8, Fp2, AF7, AF3, AFz, F1, F5, FT7, FC3, C1, C5, TP7, CP3, P1, P5, PO7, PO3, POz, PO4, PO8, P6, P2, CPz, CP4, TP8, C6, C2, FC4, FT8, F6, AF8, AF4, F2, Iz). Signals were acquired using BrainVision Recorder (BrainProducts GmbH), with a sampling rate of 1000 Hz and an online reference at FCz. Electrode impedances were kept below 10 kΩ throughout the session.

Offline preprocessing was conducted using eeglab (45) and custom scripts in MATLAB. The data were band-pass filtered between 0.5 and 45 Hz. Channels exhibiting excessive noise (average of .86 channels per participant) were removed and subsequently interpolated using spherical splines. The continuous signal was epoched with respect to the gabor onset, re-referenced to the common average and oculomotor artifacts were identified and removed using independent component analysis (ICA). Finally, trials containing residual artifacts were visually inspected and rejected (average of 35.36 trials excluded per participant).

### EEG Multivariate Pattern Analysis

To determine whether features of the visual stimulus were reliably decodable from EEG, we performed time-resolved multivariate pattern analysis (MVPA) using the Amsterdam Decoding and Modeling (ADAM) toolbox (46). The analysis focused on decoding separately spatial frequency and orientation of the stimulus from the preprocessed EEG data, aligned to stimulus onset.

For each participant, EEG epochs were down-sampled to 250 Hz and entered into a backward decoding model (BDM) with 10-fold cross-validation. A linear discriminant analysis (LDA) classifier was trained to discriminate between classes based on the scalp-wide voltage patterns at each time point. Classification performance was quantified using the area under the receiver operating characteristic curve (AUC), and decoding was performed separately at each time point from - 500 ms to 2000 ms relative to stimulus onset.

Group-level statistical inference was performed using cluster-based permutation testing (47) implemented in the ADAM toolbox. This method controls the family-wise error rate while retaining sensitivity to temporally extended effects.

### Eye-movements control analyses

Eye-tracking analyses were conducted on the subset of 19 participants with reliable data (i.e., <20% missing samples). To rule out the possibility that decoding performance during the maintenance period reflected subtle eye movements, we first performed a control analysis in which only horizontal and vertical eye-position traces were used as features in the decoding model. Second, we detected microsaccades using a velocity-based algorithm (48) and compared microsaccade rates across spatial frequency conditions. For both analyses, multiple comparisons in the temporal domain were corrected using a permutation-based cluster correction (N = 1000).

### Phase-Dependent Decoding

We introduced a novel method, phase-dependent decoding, to investigate how neural representations of visual features vary as a function of the instantaneous phase of ongoing alpha oscillations.

Continuous EEG data were first band-pass filtered in the alpha range (8–12 Hz) (or in the theta range (4-7 Hz) for a control analysis) using FieldTrip’s (49) ‘firws’ filter, a zero-phase, non-causal windowed-sinc finite impulse response filter that preserves phase information. From the filtered signal, instantaneous phase and amplitude were extracted via the Hilbert transform.

To ensure robust phase estimation and improve computational efficiency, time points with alpha amplitude below the 66th percentile were excluded. For each phase angle of interest (e.g., peak or trough), we identified all timestamps across all trials where the instantaneous alpha phase matched that angle. Around each timestamp, we extracted scalp voltage data spanning ±5% of the full phase cycle and averaged these surrounding points to enhance the signal-to-noise ratio. This yielded a set of composite voltage maps representing neural activity specifically at that phase.

Separate datasets for peak and trough phases were then created, and multivariate decoding of spatial frequency was performed using the ADAM toolbox. Decoding accuracy was then compared across peak and trough datasets with a two-tailed paired t-test. This procedure was repeated for every channel in the scalp and cluster-based permutation correction (N=1000) accounted for multiple comparisons across channels. This approach yields a spatial map of peak vs trough modulation of decoding accuracy (Figure 2B).

### Harmonic Regression

Phase-dependent decoding was computed across 18 equally spaced phase bins spanning the full oscillatory cycle (0 - 2*π*). To asses rhythmic modulation of accuracy with respect to phase, we fit a harmonic regression to the resulting phase–accuracy profiles following the methods outlined in (50). For each participant and electrode, decoding accuracy was modelled as a linear combination of sine and cosine terms:

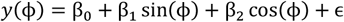

where *ϕ* denotes the phase bin centers (ranging from 0 to 2*π*), and ε is residual error. From the regression coefficients, we derived the amplitude modulation as:

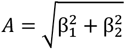

reflecting the strength of phase-dependent modulation, and the preferred phase as:

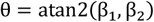

indicating the phase at which decoding accuracy peaks.

Then, we generated null distributions of amplitude and phase estimates by randomly permuting (N=1000) the phase labels and re-fitting the harmonic model. Because amplitude values are strictly non-negative, we computed robust z-scores using the median absolute deviation (MAD) of the null distribution:

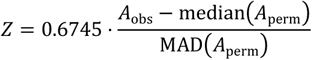

This provided a normalized, outlier-robust effect size for each participant and electrode that can be meaningfully tested against zero.

For phase, we quantified phase alignment across participants using the cosine similarity between each subject’s preferred phase and the group mean phase (i.e., the circular average):

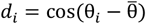

This measure is the equivalent of a dot-product in circular space and reflects phase consistency across subjects: values near 1 indicate close alignment with the group mean and values near –1 indicate opposite phase preference. We again computed robust z-scores for this alignment measure using the corresponding null distribution from the permuted data, yielding per-participant, per-channel phase alignment scores. At the group level, we tested the z-scores for both amplitude and phase alignment against zero using a one-sample t-test. To correct for multiple comparisons across electrodes, we applied a cluster-based permutation test as implemented in the FieldTrip toolbox.

### Phase-Lag Analysis

For each subject, we identified the electrodes within the significant frontal and occipito-parietal clusters (as determined by the cluster-based permutation test on peak–trough decoding differences) and computed all pairwise combinations between them (N = 243 pairs). EEG data were segmented from 1000 to 2000 ms post-stimulus and bandpass filtered in the alpha range (8–13 Hz). For each channel pair, we computed the cross-spectral density (CSD), and the phase difference was derived from the angle of the resulting complex-valued CSD. This yielded an estimate of phase differences for each channel pair and participant.

To assess whether phase differences were non-uniformly distributed and clustered around 180°, we applied a circular V-test (from the CircStat toolbox (51)) to each channel pair across participants, testing against a mean direction of 180°. The resulting p-values were Bonferroni-corrected for multiple comparisons across the 243 channel pairs.

Additionally, we computed the phase-slope index (PSI) between the same channel pairs, following the procedure described in (21). The PSI quantifies the consistency of phase differences across neighboring frequencies: when one signal systematically leads another in phase across frequencies, the PSI becomes positive, indicating a directed influence from the leading to the lagging site. Positive PSI values therefore indicate that anterior channels lead posterior ones, whereas negative values indicate the opposite direction of information flow. To assess statistical significance, a null distribution was generated by randomly sign-flipping the PSI across iterations (N=1000), and the observed value was compared against this distribution.

## Acknowledgments

This research was carried out on the High-Performance Computing resources at New York University Abu Dhabi.

This work was supported by New York University Abu Dhabi (NYUAD) Center for Brain and Health, funded by Tamkeen under NYUAD Research Institute Grant CG012.

